# Cross-species evidence for the refinement of intrinsic neural timescales supporting executive system maturation through adolescence

**DOI:** 10.64898/2025.12.12.694027

**Authors:** Shane D. McKeon, Daniel Petrie, Valerie J. Sydnor, Zhengyang Wang, Junda Zhu, Alyssa Famalette, Will Foran, Ashley C. Parr, Finnegan J. Calabro, Taylor Abel, Christos Constantinidis, Beatriz Luna

**Author notes:** Corresponding author(s): Shane D. McKeon & Beatriz Luna. Lead Contact: Shane D. McKeon.

## Abstract

Brain functional, structural, and neurochemical maturation has been found to support the specialization of executive systems through adolescence that will lead to adult level processing. Importantly, animal models and initial EEG studies in humans indicate developmental improvements in neural processing of complex information that would be evident in changes in temporal dynamics, which are not well-understood. Intrinsic neural timescales (INTs), or the temporal windows over which neural populations integrate inputs, have been proposed to reflect circuit-level properties such as excitatory-inhibitory (E/I) balance, myelination, and functional properties supporting complex information processing such as in executive functioning. Here, we used a multimodal, cross-species approach to investigate how INTs develop across adolescence to support cognitive specialization. In parallel analyses using a large longitudinal human EEG cohort, adolescent intracranial sEEG recordings, and macaque local field potentials, we observed robust reductions in INTs through adolescence, particularly in frontal and parietal association cortices. These developmental reductions were shaped by local circuit physiology, as evidenced by associations between shorter INTs and both lower aperiodic exponents, indicative of increased E/I balance, and reduced spectral offsets, suggesting lower aggregate spiking. We also found structural contributions via developmental interactions between age and deep layer intracortical myelination which predicted shorter INTs, suggesting that long-range circuitry may play a key role in shaping spontaneous temporal dynamics. Functionally, shorter INTs in adolescence were linked to improved working memory accuracy and reduced response time variability, indicating a behavioral advantage of refined temporal integration windows through development. Together, these findings establish INTs as a conserved, biologically grounded signature of adolescent brain maturation, providing a mechanistic framework for how structural and physiological refinements reorganize temporal processing to support increasingly efficient cognitive function.

## Introduction

Adolescence is characterized by improvement in cognitive abilities^1^, in parallel with specialization of brain systems that recent evidence indicate is a period of critical period (CP) in prefrontal cortex (PFC)^2–4^. Executive functions are known to show rapid improvements in childhood, with continued refinement throughout adolescence and stabilization into young adulthood^5^, suggesting a unique developmental optimization occurring during adolescence which establishes adult-like performance. While structural changes, such as synaptic pruning^6–8^ and increased myelination^9,10^ are well documented and thought to support stable, reliable, and efficient neural network function^2,11^, less is known about how population-level neural activity changes during this developmental window to support complex information processing underlying cognitive development. Animal models and initial EEG and MRI studies in humans indicate developmental improvements in neural processing of complex information including developmental decreases in neural processing^12^, increases in signal to noise ratio (SNR)^13^, enhanced effective dimensionality^14,15^, and increases in excitatory/inhibitory balance, reflecting CP^16^. These enhancements in information processing should impact temporal dynamics of neural processing, but this has not been interrogated. Here we characterize changes in temporal windows of integration, which reflect how long neural populations maintain and integrate information over time. These intrinsic neural timescales (INTs) have been shown to vary hierarchically across cortex^17^ and to relate to information processing demands, with shorter timescales supporting rapid sensory integration and long timescales supporting sustained, higher-order processing^17,18^. Developmentally, a shift toward shorter, more efficient timescales may reflect increased temporal precision and flexibility as cortical circuits mature. Given known increases in myelination^9,10,19^, enhancements in information processing and increases in E/I balance^16,20^ that should expedite complex information processing mechanisms^13–15^, we hypothesize that intrinsic neural timescales will shorten across adolescence, reflecting a refinement in temporal processing that underlies improvements in executive function. Moreover, we predict that this shortening is supported by increases in excitation/inhibition balance and cortical myelination. In this work, we test these hypotheses using encephalogram (EEG) recordings, intracranial stereo-EGG (sEEG) recordings, macaque local field potentials (LFPs), and myelin derived from quantitative T1 maps, to link neuroelectrical and microstructural changes to developmental refinements in neural dynamics and cognitive control.

Patterns of neural activity are temporally dynamic, allowing the brain to integrate incoming information across various time windows, tailoring the response to changing environmental demands^21^. These time windows of integration are known as intrinsic neural timescales (INT), defined as the length of time in which incoming information can affect a response^22^. Intrinsic neural timescales can be interrogated via the decay properties of the autocorrelation function (ACF)^17,22–26^, and reflect how neural activity at one moment relates to activity at future time points. The autocorrelation function captures the consistency of these temporal relationships, where the decay indicates how quickly a system loses memory of its prior state. Across the cortical hierarchy, shorter vs longer timescales can be advantageous in different regions. A shorter INT may promote temporal segregation, allowing the system to distinguish rapid progressions of stimuli from one another with precise temporal encoding, leading to multiple short evoked activity profiles lasting the duration of the incoming stimuli^22^. In contrast, longer INTs lead to correlated neural activity across time, allowing multiple inputs to summate and produce a longer evoked activity^22^. Sensory unimodal regions need to respond rapidly to incoming stimuli, and therefore tend to have shorter timescales compared to higher-order transmodal areas, which integrate and accumulate information more slowly^17,27–30^. On the microscale, spiking activity at the neuronal level can influence the temporal structure observed at the macroscale, such as in EEG. Regions with more persistent or recurrent spiking show longer INTs, suggesting a neuronal mechanism for prolonged temporal integration^21,31^.

Within higher-order regions, such as the prefrontal cortex (PFC), INTs can also be dynamically modulated depending on the task demands, with longer timescales apparent during cognitive processing relative to resting state^27^. For example, during working memory (WM) tasks, the PFC exhibits shorter INTs during encoding and longer INTs during WM maintenance^32,33^, suggesting that INTs are functionally dynamic and can modulate their length depending on cognitive demands. These dynamic timescales may depend on spatially-local connectivity, as computational modeling has suggested slow timescales increase with the strength of recurrent interactions^33,34^, with shifts of recurrent spatial connectivity and fluctuations in network excitability potentially supporting task-dependent INT modulation^33^. Task-dependent modulation of neural timescales appears to be mediated in part by N-methyl-D-aspartate (NMDA) receptor-dependent recurrent excitation, as well as fast acting inhibitory mechanisms, such as those mediated by GABAergic interneurons^32^. Furthermore, spatial gradients in neuronal timescales across the cortex have been found to correlate with the regional expression of genes involved in excitatory and inhibitory signaling. These include genes encoding NMDA receptor subunits, GABA-A receptors, and markers of specific inhibitory cell types such as parvalbumin-expressing (PV+) interneurons^32^.

Computational modeling has also shown increases in recurrent excitation or decreases in inhibition, leading to increased E/I balance, and prolonged INTs^28^. This suggests that regional changes in the E/I balance may directly support the emergence of functional timescale hierarchies across the brain, linking molecular architecture to circuit-level temporal processing. Developmentally, previous work has shown that the E/I balance increases across adolescence^2,16,20,35–38^, driven by the maturation of inhibitory circuitry, which in turn suppresses spontaneous oscillatory, asynchronous activity, in favor of evoked synchronous activity^13,39,40^. Importantly, here we distinguish E/I balance, reflecting the coordination of excitatory and inhibitory activity to stabilize network function, from the E/I ratio, which typically decreases with development as inhibition strengthens, and excitation dampens. The PFC itself continues to mature, relying increasingly on local gain control and balanced excitation-inhibition (E/I) dynamics to integrate these inputs over shorter timescales^16,20,41,42^. Furthermore, recent work has linked temporal dynamics to underlying neurobiological mechanisms such as synaptic pruning^23,43^ and myelination^32,33^. More specifically, synaptic pruning, known to continue to occur in the PFC during adolescence^6,8^, has been suggested to reduce recurrent excitation, leading to more transient neural responses and ultimately shorter INTs^24,44^.

While prior work has well-characterized regional differences in INTs in adults, less is known about how these timescales evolve during adolescence. Importantly, while previous studies have examined developmental decreases in spontaneous brain activity^13^, how these age-related decreases translate to changes in temporal integration, specifically during task demands, remains mechanistically unclear. Thus, the present study addresses these gaps by investigating developmental changes in INTs across cortical regions through adolescence into adulthood. Further, direct evidence linking macro-scale population-level INTs to microscale neural spiking patterns remains limited, particularly in humans where invasive recording is rare due to ethical and practical constraints. To bridge this gap, the present study includes local field potentials (LFPs) recorded from human intracranial stereo-EGG (sEEG) from pediatric epilepsy patients, as well as a cross-species examination leveraging recordings from adolescent non-human primates (macaques). These unique data offer improved spatial and temporal resolution compared to scalp EEG and provide a closer approximation to the underlying neuronal dynamics and mechanisms supporting the development of INTs through adolescence and potentially provide confirmatory evidence for scalp EEG findings. By examining INTs across EEG, sEEG, and macaque LFPs we aim to clarify how developmental changes in population-level temporal dynamics emerge from local circuitry and whether such temporal tuning is conserved across species. Based on these data, we sought to examine the developmental trajectory of task-dependent INT modulation and its relationship to developmental trajectories of executive function. Furthermore, given evidence that the E/I balance and myelination have been shown to shape the temporal structure of neural activity^32,36,45^, we tested whether INT development across the cortex was associated with the developmental increases in EEG-derived measures reflecting E/I balance, and increases in intracortical myelin, measured using ultra-high field quantitative MRI. Finally, to test whether population-level temporal dynamics were reflected in local circuit dynamics and whether developmental changes in INTs were conserved across species, we examined the maturation of INTs across both sEEG and macaque LFP data. These data support a model of adolescent development in which maturation of circuit level neural properties allows for dynamic fine-tuning of temporal properties, supporting the emergence of reliable, adult-like cognitive processing.

## Results

We studied intrinsic neural timescales via the EEG-based autocorrelation window (ACW) across frontal, parietal, and occipital cortices in an accelerated longitudinal sample of 161 individuals 10-32 years old with 1-3 timepoints 18 months apart each, resulting in a total 290 sessions. We quantified the ACW as a measure of intrinsic neural timescale, defined as the time lag at which the autocorrelation function (ACF) decays to 50% of its maximum. This value, referred to as ACW-50 (here referred to as “ACW” for simplicity), reflects the temporal “half-life” of a signal’s self-similarity and has been established in prior work^21,24^. Given prior work suggesting that intrinsic neural timescales (INTs) are modulated by task state, we obtained ACWs for 4 minutes rest and during a memory guided saccade working memory task (MGS). Given that we did not find significant differences in ACWs between the delay and fixation epochs of the MGS task in frontal, parietal, or occipital regions (all p > 0.15), we averaged delay and fixation values within each region to derive a unified cognitive task-state measure to compare with rest.

### Intrinsic neural timescales decrease across adolescence

A central aim of this work was to assess the hypothesis that intrinsic neural timescales, as indexed by the ACW, decrease across adolescence in both rest and cognitive state, across frontal, parietal, and occipital regions. To assess this hypothesis, we first employed individual semiparametric generalized additive mixed models (GAMMs) to ACW within each region, covarying for cognitive state (rest vs task). We found significant developmental decreases in the ACW across adolescence in the frontal (edf = 1.79, *F* = 66.78, *p <* 0.001; Figure 1A), parietal (edf = 1.67, *F* = 64.30, *p <* 0.001; Figure 1A), and occipital (edf = 1.75, *F* = 47.69, *p <* 0.001; Figure 1A) regions, indicating that intrinsic neural timescales shorten across adolescence. Furthermore, we found cognitive state to be a significant moderator of ACW in the parietal (β = 0.0007, t = 2.94, *p* = 0.003) and occipital (β = 0.001, t= 4.62, *p* < 0.001) regions, with higher ACW values during task relative to rest. We then tested whether age-related changes in INTs differ between task and rest, using GAMMs with an age-by-task interaction. In the frontal cortex, we found no significant age-by-task interaction (edf = 1.45, *F* = 3.56, *p* = 0.10). However, in the parietal (edf = 1.86, *F* = 10.33, *p* = 0.001) and occipital (edf = 1.89, *F* = 8.33, *p* = 0.01) cortices we find significant age-by-task interactions, suggesting the developmental trajectory differs by cognitive state, with age-related decreases more pronounced in resting state compared to task state.

**Figure 1.**
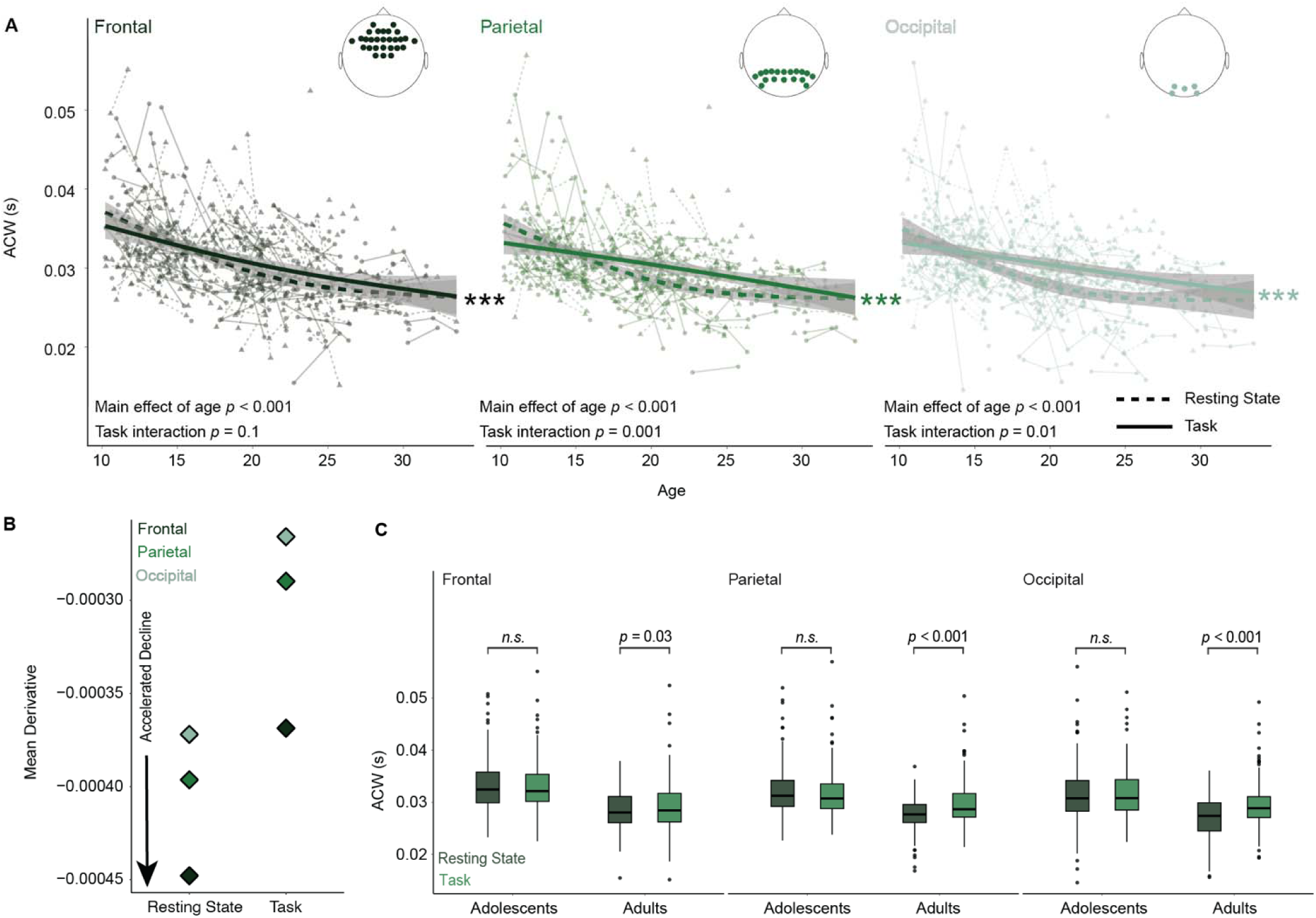
Intrinsic neural timescales decrease through adolescence to adulthood in rest and task states. A) Autocorrelation window as a function of age in frontal, parietal, and occipital regions, for resting state (dashed line) and the cognitive task state (solid line). Significant differences in age functions between task and rest are observed in all regions. Brain plots represent electrodes corresponding to each region. B) The rate of ACW decline with age accelerates from occipital to frontal regions, highlighting more pronounced developmental reductions in frontal areas. C) Boxplots compare the autocorrelation window between resting state and task state for adolescents and adults across regions, with significant differences in the frontal (p = 0.03), parietal (p<0.001) and occipital (p<0.001) regions, for adults only, as assessed with paired-samples t-tests. This suggests the differences seen between the task and rest are driven by the adult population.

We next quantified the rate of age-related decline in INTs across regions by computing the average first derivative of the age-related smooth function from each GAMM, with a more negative average derivative reflecting a steeper maturational decline. Separate GAMMs were estimated for each region and task state (rest and cognitive state). Across both states, the rate of decline was greatest in the frontal cortex, followed by the parietal, with the occipital cortex showing the shallowest decline, indicative of an anterior-posterior gradient in adolescent developmental change (Figure 1B). Notably, the overall magnitude of decline was larger during rest compared to cognitive task, in all lobes, suggesting that resting-state INTs may more directly reflect developmental changes in underlying neurobiology, such as increasing myelination or maturation of E/I balance. This may also reflect that the cognitive task requires an established pattern of processing compared to rest, which may reduce inter-individual variability in timescales, thereby attenuating age-related differences for cognitive task relative to rest.

To assess how cognitive task vs resting state differentiates across age, we averaged ACW values in each region for adolescents (< 18 years old) and adults (>= 18 years old). In contrast to adolescents, adults showed significant state-dependent differences in ACW. Paired *t*-tests revealed higher ACW values during task compared to rest in the frontal (t = 2.10, *p* = 0.037; Figure 1C), parietal (t = 5.07, *p* < 0.00; Figure 1C), and occipital regions (t = 6.13, *p* < 0.001; Figure 1C) in adults only. These results suggest that intrinsic timescales are more state-sensitive in adulthood, with longer ACWs during task engagement relative to resting state, potentially suggesting that adults have a greater ability to tune ACWs based on task demands.

### Shorter timescales reflect E/I- driven neural dynamics

Given evidence that E/I balance results in shorter intrinsic neural timescales^45–48^, and our previous EEG study looking at the developmental changes in the aperiodic exponent reflecting increases E/I balance^16^, we next investigated the association between age related changes in the aperiodic exponent and intrinsic neural timescales. The aperiodic exponent has been previously linked with changes in the E/I balance, supported by computational^49,50^, and pharmacological^51–53^ work, as well as our own empirical studies linking it to 7T MRSI-derived measures of GABA and Glutamate^16^. Derived from the slope of the power spectral density (Figure 2A), flatter slopes were associated with increases in E/I balance, which increased from adolescence to adulthood (Figure 2A). Thus, we hypothesized that shorter intrinsic neural timescales would be associated with a lower aperiodic exponent (i.e. a flatter slope). We quantified the aperiodic exponent at each electrode and averaged values across frontal, parietal, and occipital regions, consistent with our approach for ACW. We first examined correlations between ACW and aperiodic exponent, finding strong positive associations in all three regions (frontal: *r* = 0.61; parietal: *r* = 0.64; occipital: *r* = 0.74) reflecting that flattening of the exponent corresponded with shorter ACWs. Linear models that controlled for age further confirmed these associations, with flattening of the aperiodic exponent being significantly being associated with shorter ACW in the frontal (β = 28.8, *t* = 10.4, *p* < 0.001), parietal (β = 30.86, *t* = 11.15, *p* < 0.001), and occipital (β = 45.65, *t* = 16.81, *p* < 0.001) regions (Figure 2B). These results suggest that INTs may be modulated by underlying shifts in E/I balance, providing empirical support for a mechanistic link between local circuit dynamics and macroscale temporal processing, via EEG.

**Figure 2.**
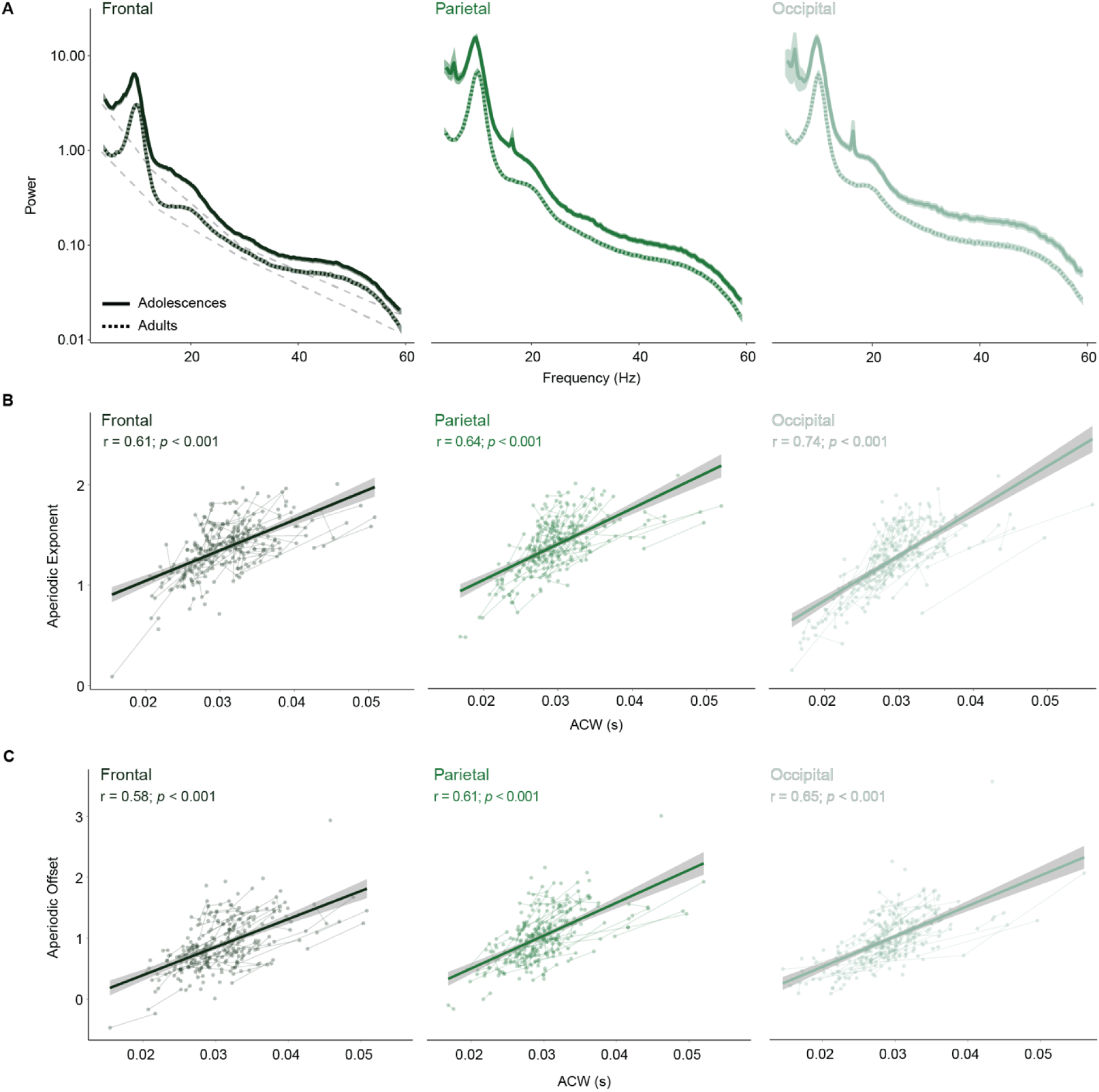
Shorter intrinsic neural timescales are associated with EEG-derived measures of E/I balance and aggregate spiking activity. A) Power spectra in frontal, parietal, and occipital regions from adolescents (solid lines) compared to adults (dashed lines). The aperiodic exponent was derived from the slope of the 1/f component of the power spectral density (illustrated by the light gray dotted lines in the frontal region). This exponent reflects the rate at which power decreases with increasing frequency. The aperiodic offset, derived from the y-intercept, reflect neural aggregate spiking. B) Shorter autocorrelation windows (ACW) are associated with a lower aperiodic exponent, in the frontal (left), parietal (middle), and occipital (right) lobes, indicative of more balanced excitatory and inhibitory neural dynamics. C) ACW is also positively associated with aperiodic offset in frontal (left), parietal (middle), and occipital (right) lobes, suggesting that reductions in aggregate neural spiking (indexed by offset) may be underlying shorter intrinsic neural timescales.

The aperiodic component of the EEG signal is also described by another metric, derived from the amplitude of the power spectrum, known as the aperiodic offset. The offset reflects broadband shifts in power^54^, which are thought to reflect the magnitude of neuronal population spiking^54,55^. Thus, we hypothesized that more aggregate spiking, reflecting persistent firing and longer temporal integration^17,21,25^, would be associated with a longer ACW. To assess the relationship between offset and ACW we first examined correlations, finding strong positive associations in all three regions (frontal: *r* = 0.58; parietal: *r* = 0.61; occipital *r* = 0.64) indicating that less aggregate spiking corresponded to shorter ACWs. We once again used linear models to further investigate these associations while controlling for age, and observed a similar patter with lower aperiodic offset was significantly associated with shorter ACW in frontal (β = 34.52, *t* = 10.14, *p* < 0.001), parietal (β = 40.40, *t* = 12.32, *p* < 0.001), and occipital (β = 37.19, *t* = 13.91, *p* < 0.001) regions (Figure 2C). These results suggest that INTs may be impacted by population level aggregate neuronal activity, such that greater activity contributes to prolonged windows of temporal integration.

### Shorter timescales support working memory performance

Given that ACW values are modulated by task demands, we investigated how developmental decreases in ACW, observed in both rest and task, were associated with developmental improvements in working memory task performance, reflected in increases in accuracy and decreases in response latency^12,16^. We first assessed the main effect of across-state (rest and task) ACW on accuracy (measured as the distance in degrees of visual angle of the memory guided eye movement from the intended target) in each region. Here, we found significant positive associations in the frontal (β = 18.40, t = 2.41, *p =* 0.048; Figure 3A) and parietal (β = 21.53, t = 2.62, *p =* 0.027; Figure 3A) regions, indicate that lower ACWs were associated with better task performance (fewer degrees from target), but not in the occipital region, highlighting the cognitive, rather than sensory, aspect of this measure (β = 18.06, t = 2.36, *p =* 0.055; Figure 3A). No main effects of cognitive state (task vs. rest) on task performance were observed, indicating an individual person specific relationship to ACW to cognitive performance, as opposed to a task dependent one. In contrast, trial-by-trial variability of accuracy was not significant associated with ACW in any region (all *p >* 0.05; Figure 3B). For response latency, there were no significant associations with ACW in any region (all *p >* 0.05; Figure 3C); however, trial-by-trial variability in response latency had a significant positive association in frontal (t = 3.244, *p =* 0.004; Figure 3D), but not in parietal (t = 1.81, *p =* 0.21; Figure 3D) nor occipital (t = 1.32, *p =* 0.55; Figure 3D), indicating that lower frontal ACWs were associated with decreased trial-by-trial variability. In sum, faster INTs were associated with higher working memory accuracy (in frontal and parietal regions) and lower response time variability (frontal only), yet not with accuracy variability or average response latency.

**Figure 3.**
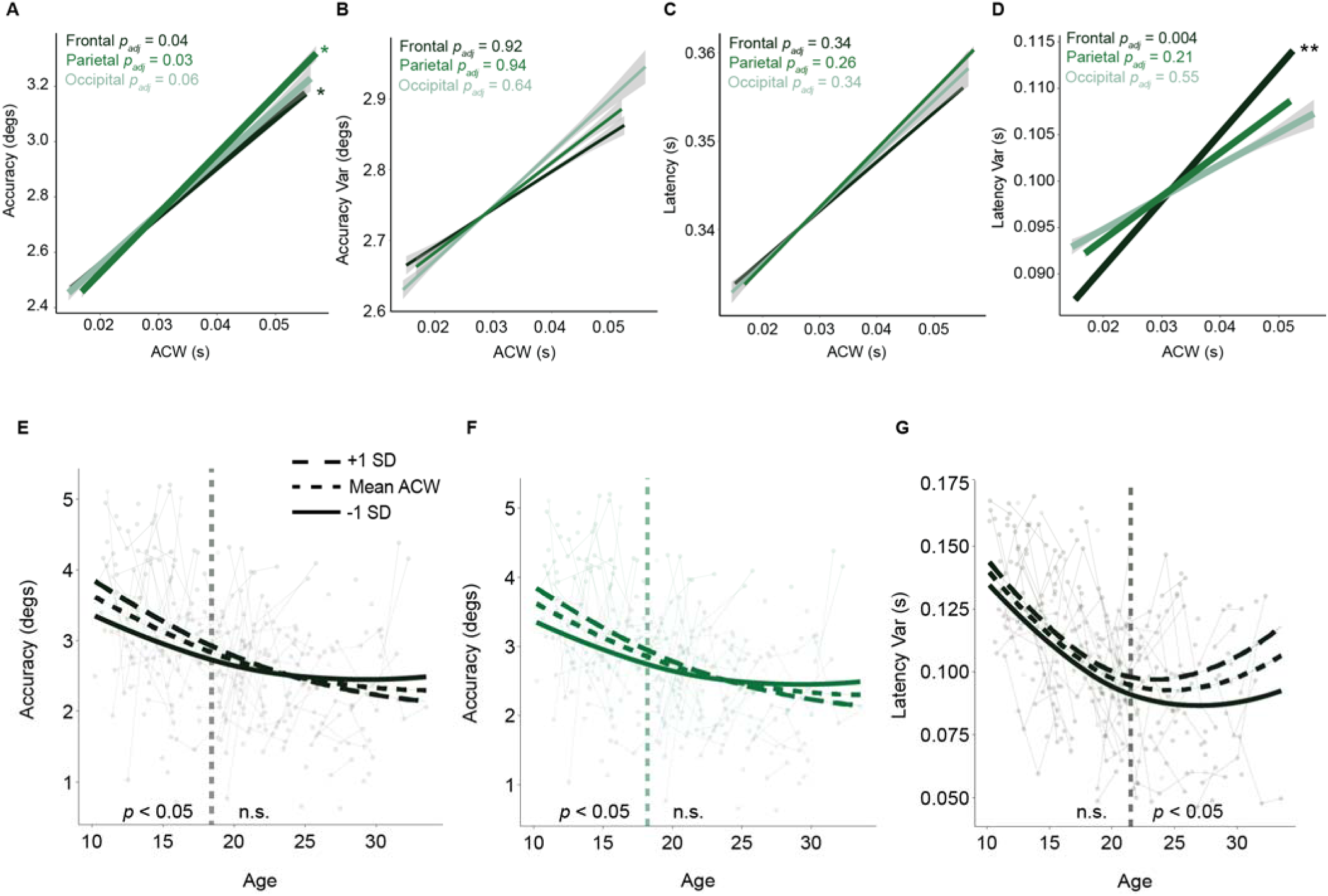
Shorter intrinsic neural timescales are associated with improved working memory performance. A) Accuracy (measured as the degrees of visual angle away from the intended target) improves with shorter autocorrelation windows in the frontal and parietal lobes. B) Accuracy variability (measured as the standard deviation across trials) does not change with ACW in any region. C) Response latency does not significantly change across the ACW in any region. D) Latency variability (measured as the standard deviation across trials) decreases with shorter autocorrelation windows in the frontal lobe. E-G) Age-varying relationships between ACW and performance, in which the dotted line represents the average ACW across participants, the dashed line represents one standard deviation above the mean, and the solid line represents one standard deviation below the mean. Vertical dashed lines mark age boundaries where effects reach statistical significance (p < 0.05). E) The effect of long (+1 SD), average, or short (-1 SD) ACWs on MGS accuracy across age in the frontal region. Differences in ACW had a significant effect on accuracy under 18.4 years of age, with shorter ACWs being advantageous in adolescence. F) The effect of long (+1 SD), average, or short (-1 SD) ACWs on MGS accuracy across age in the parietal region. Differences in ACW had a significant effect on accuracy under 18.2 years of age, with shorter ACWs being advantageous in adolescence. G) The effect of long (+1 SD), average, or short (-1 SD) ACWs on MGS latency variability across age in the frontal region. Differences in ACW had a significant effect on latency variability over 21.5 years of age, with shorter ACWs being advantageous in adulthood.

To examine age-dependent relationships between ACW and behavioral performance, we modeled the age-varying effect of ACW using a GAMM for accuracy and variability in response latency. This approach allows the strength of the ACW–behavior relationship to vary continuously with age^56^. We applied this method to the significant associations described above. For accuracy in the frontal region, this model revealed a significant interaction between age and ACW (F = 5.58, *p =* 0.004), indicating that the influence of ACW on behavioral accuracy differed across development. A comparison of the first derivative of the age term from the GAMM model, where the confidence interval indicates ages where the ACW was significantly related to accuracy, reflected a significant association between ACW and accuracy only prior to 18.4 years of age, with shorter ACWs being advantageous in adolescence (Figure 3C), potentially reflecting more advanced maturation. A parallel analysis in the parietal cortex yielded similar results, with a significant age-by-ACW interaction (F = 4.34, *p* = 0.013) and a comparable developmental window below 18.2 years in which shorter ACWs were again associated with better performance (Figure 3D). Based on our previous significant ACW-behavioral associations, we further interrogated the relationship between latency variability and ACW in the frontal cortex. Here, we found a significant age-by-ACW interaction (F = 18.38, *p <* 0.001) and a significant developmental window *above* 21.5 years where shorter ACWs were associated with less response latency variability only in adults (Figure 3E). These results suggest that the behavioral relevance of INTs is not fixed across development but instead shifts, with shorter INTs supporting enhanced task accuracy in adolescence yet more stable, efficient responding in adulthood.

### Shorter timescales are supported by cortical myelination

During adolescence, myelination within the gray matter continues across the cortex, with sensorimotor regions showing greater overall increases, while association areas such as the prefrontal cortex exhibit more protracted, though less extensive, myelination^10,57–61^. Previous studies have suggested that myelin may play an important role in shaping the temporal dynamics of neuronal activity^10,62^, such as in signal timing and synchronization. Additionally, previous studies have explored the connection between intrinsic neural timescales and myelin content in adulthood, finding shorter timescales in more heavily myelinated regions^32^. Thus, we hypothesized that developmental increases in myelination, as we have previously reported in the same participants^10^, would be associated with the decreases in ACW reported here. To test this, we examined the relationship between ACW and intracortical myelination. Myelination was indexed by R1, derived from quantitative MP2RAGE scans collected at 7T MRI, at cortical vertices corresponding to the EEG electrode locations when projected onto the surface using a surface-based atlas (Figure 4A; see Methods). R1 was then averaged into superficial depths (between the 20th and 40th percentile of cortical thickness) and deep depths (50 to 80 percentile of cortical thickness) in regions corresponding to each electrode label. We then related ACW values from both rest and task states in the frontal, parietal, and occipital cortices to these vertex-wise R1 measures. Neural activity from both deep and superficial cortical layers contribute to scalp-recorded EEG activity^63,64^, thus, we fit independent GAMMs to study the relationship between ACW (rest and task) and both deep and superficial R1. Individual GAMMs were fit for each region. While we did not find any significant main effects of ACW (task nor rest) on R1 (deep nor superficial) for any of the regions (Figure ***4***B), we did find significant ACW-by-depth interactions in frontal (F = 6.83, *p =* 0.02) and parietal (F = 6.16, *p =* 0.03) regions during resting state for deep R1 (Figure ***4***C). Although the main effect of ACW was not significant across the full sample, partial smooths revealed significant age-related patterns within specific depths for both frontal and parietal, indicating that developmental trajectories of ACW differ meaningfully by cortical depth, where deep depth myelination, which supports top-down connectivity may more strongly influence temporal dynamics of neuronal activity.

**Figure 4.**
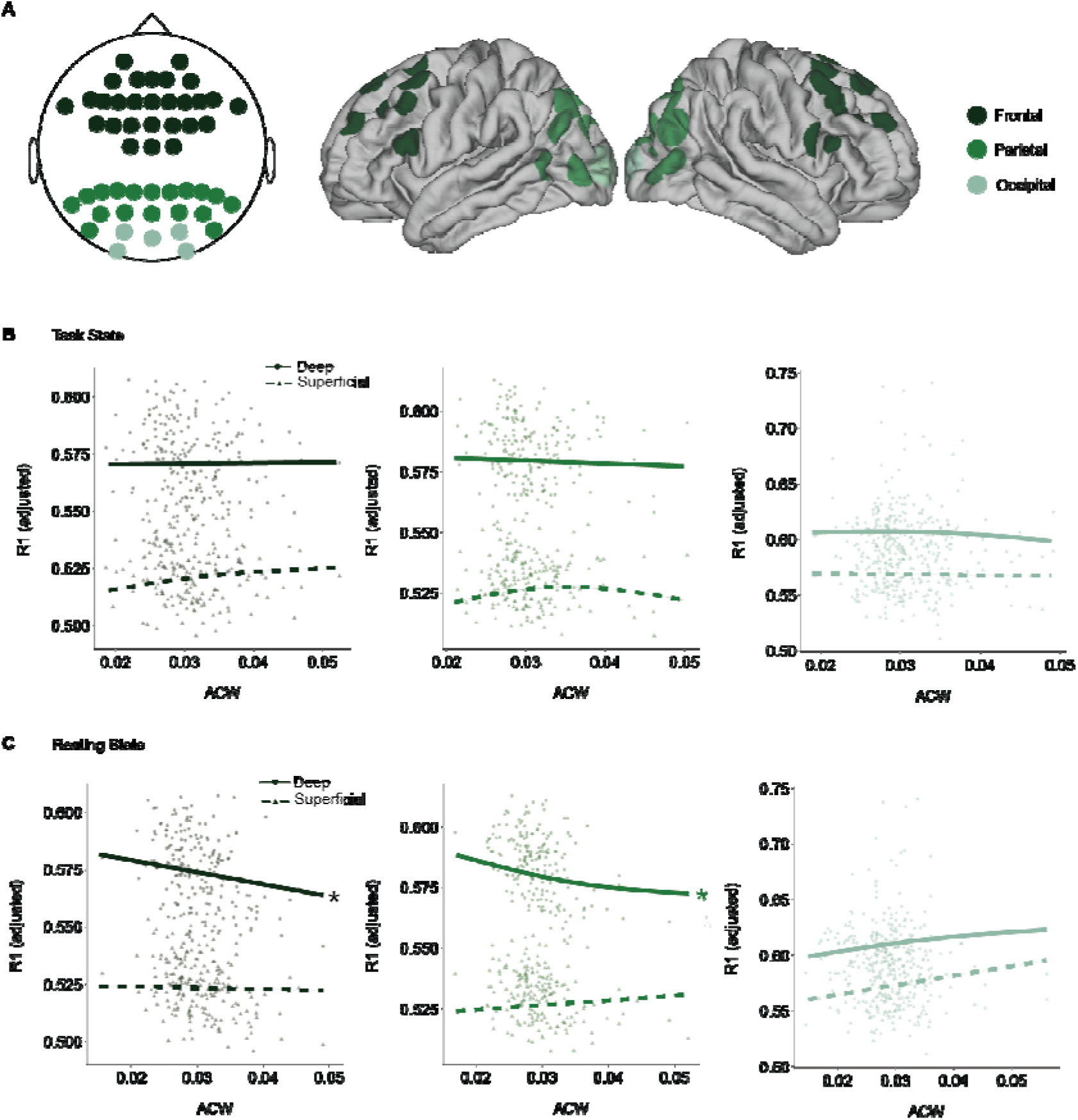
Intrinsic neural timescales are associated with R1 in the frontal and parietal regions during rest. A) The positions of EEG electrodes on the scalp were mapped to corresponding locations on the cortex to create a surface-based atlas. Average depth-dependent R1 and the ACW values were calculated by averaging measurements from spatially proximate electrodes within the frontal, parietal, and occipital cortices. B) R1- task state ACW associations are shown for deep and superficial layers in the frontal (left), parietal (middle) and occipital cortex (right); data points represent partial residuals from a depth interaction model that controlled for age. C) R1- resting state ACW associations are shown for deep and superficial layers in the frontal (left), parietal (middle) and occipital cortex (right); data points represent partial residuals from a depth interaction model that controlled for age. GAMMs with a depth interaction term reveal a significantly stronger negative association between R1 and the resting state ACW in deep as compared to superficial cortex, in the frontal (left) and parietal (middle) cortices.

### Developmental reduction in timescale replicate in human sEEG

To validate the developmental changes in intrinsic neural timescales observed in our EEG analyses, we investigated age-related changes in the ACW using intracranial recordings in humans. Specifically, we quantified the ACW in resting state intracranial stereo-EEG (sEEG) data from a separate developmental dataset of 14 medication-resistant epilepsy patients ages 9-21 years old (representation of electrode placement in Figure 5A). A complete table of patient demographics and number of electrode placements can be found in Supplemental table 1. This complementary dataset provides direct recordings of neural activity from cortical sources with high spatial and temporal precision, enabling us to assess whether the age-related shortening of ACW observed in scalp EEG generalizes across recording modalities.

**Figure 5.**
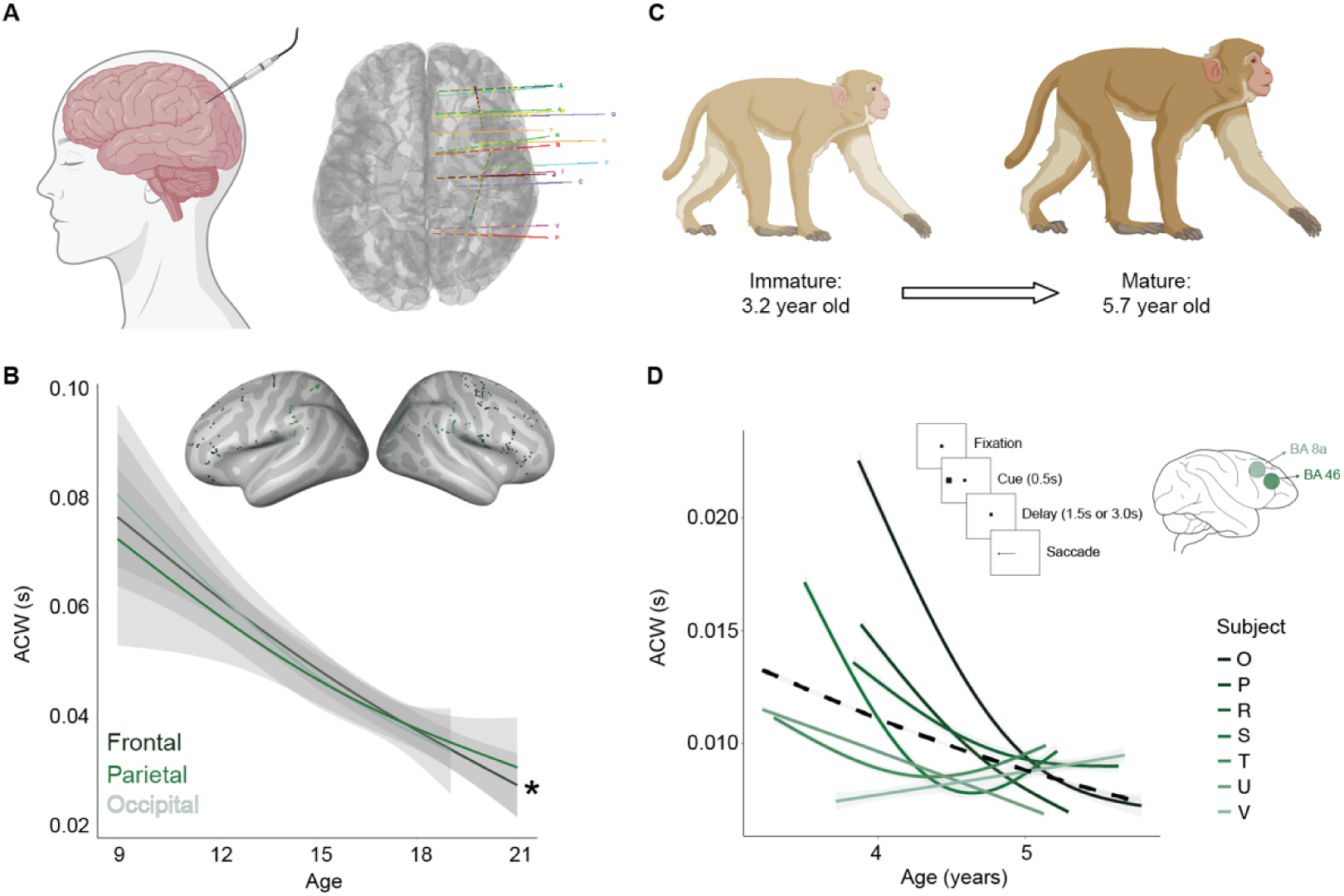
Developmental decreases in intrinsic neural timescales replicate in human sEEG and macaque LFPs. A) Schematic representing sEEG electrode (left) and representative scan and electrode placement derived from brainstorm (right). B) GAMM smoothed developmental trajectories of the resting state ACW in the frontal, parietal, and occipital cortices, showing significant decreases across adolescence in the frontal cortex. Brain plots show non-seizure electrode placements across all subjects, colored according to lobe, frontal, parietal, or occipital. C) Schematic representing the non-human primate macaque dataset, ranging from ages 3.2-5.7 years of age. D) GAMM smoothed developmental trajectories of the ACW for each macaque (greens). A group-level trend line was overlaid using a GAM with an unfixed smoothing parameter (dashed line), capturing the overall developmental pattern across the cohort. Macaque data was derived from the fixation and delay period of the ODR task (upper right corner, lefthand side) and recorded from two brain areas: BA 8a and BA 46 (upper right corner, righthand side).

Non seizure labeled electrodes from the sEEG dataset were grouped across participants and classified into frontal, parietal, and occipital regions. The number of non-seizure electrodes per subject are designated in supplemental table 1. Using GAMMs to study the smoothed developmental trajectories of the resting state ACW in the frontal, parietal, and occipital cortices, we found significant age related decreases in the frontal (F = 6.73, *p* = 0.036; Figure 5B) cortex, but not in parietal (F = 4.88, *p =* 0.06; Figure 5B) nor occipital regions (F = 0.865, *p =* 0.74). These results provide strong support that the developmental trajectories identified in our EEG dataset (Figure 1) reflect fundamental changes in local neural circuit dynamics.

### Macaque LFPs replicate developmental reductions in timescales

To further characterize the neural basis of ACW changes, we extended these analyses to non-human primates, we analyzed local field potentials (LFPs) recorded from 7 awake macaque monkeys ages 3.2-5.7 years old (average age of puberty onset is around 3.3 years^65^). LFPs were recorded from BA 8a and 46, corresponding to the dorsolateral prefrontal cortex (DLPFC), during an Oculomotor Delay Response (ODR) task similar to the MGS task employed in our scalp EEG study (Figure 5D upper right corner), at intervals of approximately three months. ACW was calculated on each channel, for each behavioral session, for every trial. Developmental trajectories were modeled using one GAMM with random slopes for each macaque. By leveraging this independent dataset, we aimed to test the consistency and spatial specificity of ACW developmental trajectories and their potential conservation across primate brains.

Here, we found significant decreases in ACW across adolescence during working memory (*p* < 0.0001; Figure 5D). These findings provide evidence that INTs decrease with age in the macaque prefrontal cortex, mirroring the developmental trajectories observed in our neurotypical EEG dataset. This cross- species convergence suggests that INT maturation is conserved across development in the primate lineage.

## Discussion

The current study demonstrates that intrinsic neural timescales (INTs), as measured by the autocorrelation window (ACW), consistently decrease across adolescence within the frontal and parietal lobes and are modulated by task state in adulthood. Using a multimodal, cross-species approach, including longitudinal developmental human scalp EEG, adolescent human sEEG, and macaque LFP recordings, we provide evidence that developmental refinement of temporal dynamics may reflect cortical specialization that is shaped by underlying changes in E/I balance and intracortical myelination. Specifically, we found INTs to decrease across adolescence, a phenomenon that was upheld regardless of modality (scalp vs intracortical recordings) as well as across species, suggesting a fundamental biological process of the shortening of temporal windows of integration across adolescence. Consistent with previous literature^32,33^, we also found INTs to be modulated by cognitive state, with increased windows of integration occurring during task state vs resting state but only in adults. By combining our INT analyses with our previous aperiodic EEG analysis, we uncovered that a lower aperiodic exponent and a lower offset, indicative of increased E/I balance and lower neural aggregate spiking, respectively^54,55^, are associated with shorter INTs. We then aimed to investigate the underlying mechanisms supporting the reduction in INTs by associating our EEG measures of INTs with a quantitative MRI measure of intracortical myelin. Here, we found deep myelination may influence temporal dynamics of neuronal activity as opposed to superficial. Finally, using a working memory saccade task, we show that shorter INTs were associated with improved working memory performance, furthering demonstrating a role for temporal tuning in neuronal populations in cognitive development. Together, these results suggest that the shortening of INTs is a biologically grounded index of adolescent brain development, where temporal dynamics undergo targeted refinement via intracortical myelination and E/I balance that support the emergence of mature cognitive functions.

INTs are thought to be the time window over which brain regions integrate inputs and are hypothesized to be the result of numerous neuronal mechanisms, such as synaptic efficiency, reduced temporal redundancy, pruning, myelination, E/I balance, and/or recurrent connectivity^32,66^. Adolescence is a time period where such mechanisms undergo significant maturation, including, but not limited to, increases in synaptic pruning^6–8^, increased myelination^9,10^, increased connectivity to the prefrontal cortex^67,68^, and increased E/I balance^16,20,38^, all of which have been shown to impact executive function^2,10,16,69–71^. Furthermore, prior studies have shown INTs vary across cortical hierarchies, with association regions typically exhibiting longer timescales than sensory regions in adults^17,27^. In complement to these findings, INTs are modulated by task-states, with longer timescales occurring during executive function compared to resting state^27^ in adults but not in adolescents. However, little is known about how these temporal integration windows develop during adolescence, and the extent to which they impact cognitive function. Our first key finding addresses this gap, demonstrating that INTs systemically decrease across adolescence, particularly in the frontal and parietal regions. Specifically, the frontal cortex showed the highest rate of decline across adolescence and more pronounced task modulation, reinforcing its central role in both adolescent neurodevelopment and task-based integration. We show this developmental decline in INTs across scalp EEG in a large longitudinal sample of neurotypical adolescents, replicate this finding in adolescent intracranial sEEG data, and extend to macaque LFPs, highlighting the robustness of this phenomenon across modalities and species. Functionally, this shift may reflect a developmental optimization of neural dynamics where shorter INTs, as measured by shorter windows of temporal autocorrelation, allow for more rapid updating of information, an enhanced ability to process inputs with finer granularity, and a more focused temporal integration^72^.

The neural basis of these developmental findings is further contextualized by evidence from non-human primates. In macaques, our results from local field potentials (LFPs) yielded similar developmental reductions in INTs. However, previous work quantifying timescales in the same macaques using recorded spiking activity from the same regions did not show any developmental change in neuron-specific timescales^15^. This dissociation implies that the observed reduction in INTs across adolescence may not reflect changes at the level of individual neurons but instead arise from shifts in local populations of neurons. LFPs, like EEG, capture aggregate synaptic and dendritic activity from networks of neurons, which are shaped by numerous factors, such as activity synchronization^13,39,40^, E/I balance^16,20,38^, and recurrent circuitry^67,68^, all of which significantly mature during adolescence. Similarly, intracranial sEEG recordings, which capture activity from networks of neurons, demonstrate the same developmental findings during rest, reinforcing the idea that developmental reductions in INTs are capturing population-level temporal processing rather than isolated neuronal firing. Together, these cross-species and cross-scale results suggest that developmental changes in INTs reflect a reorganization of population level circuit dynamics that is present both in rest and task.

Hierarchical INT development may reflect specialization of each region for behaviorally relevant computations, where sensory cortices must encode rapid sensory inputs, while association cortices must integrate long-term information^22,27,32,33,66,73^. These dynamic intrinsic timescales may emerge from local patterns of spatial connectivity, where prior studies have shown the organization of timescales correlates with the gradients in the strength of neural connections^33^. While maintaining the same spatial pattern, timescales have been found to be modulated by cognitive state^18,21,31–33,73^. Computational modeling has shown that timescales emerge from spatiotemporal population dynamics, shaped by local and long-range connectivity, but can be modulated by increases in the efficacy of recurrent interactions as would be required for cognitive processing^33^. Specifically, longer intrinsic timescales have been associated with increased functional connectivity, particularly in higher-order association cortices, suggesting that prolonged temporal integration may support broader network integration^17,24,32,33,74^. For example, during working memory tasks, the PFC exhibits shorter INTs during stimulus encoding but longer INTs during the greater computational requirement of maintenance, compared to rest, reflecting a functional modulation of temporal integration depending on task demands^32,33^. This suggests that INTs are not fixed but dynamically regulate to support increased computational requirements of cognitive processing. Our results of increased INTs during a working memory “task state”, when sensory input, maintenance, and motor planning must be coordinated, compared to resting state provide evidence for this process. Interestingly, the increase in intrinsic timescales from rest to task was observed only in adults (ages 18 and older), suggesting a maturational shift in when temporal receptive windows may lengthen to support sustained cognitive processing. This shift has been proposed to be mediated by the maturation of local circuit properties^6–10,67,68^, including excitatory properties such as NMDA receptor subunit composition shifts from NR2B to NR2A^32,75,76^, resulting in faster and more refined responses, as well as an overall decrease in NMDA receptor density leading to less excitation. Thus, in adolescence decreased E/I balance may result in persistent larger timescale processing regardless of state requirements. While this shift reflects a decrease in temporal integration capacity at the circuit level, creating more transient responses, it may also signal a transition toward more efficient, flexible processing as the brain matures, supporting faster information updating and more dynamic network interactions. In conjunction, inhibitory interneurons, such as PV+ cells, mediate fast, phasic inhibition that matures throughout adolescence^36,77–81^, leading to faster, more temporally precise inhibition resulting in short INTs. Taken together, reductions in NMDA- mediated excitation and increases in GABAergic inhibition lead to increased E/I balance that supports faster more temporally precise computations, such as those necessary for rapid sensory processing and cognitive flexibility, which may be reflected here as shorter intrinsic neural timescales.

To further probe the contributions of the E/I balance and the physiological mechanisms underlying the developmental reductions in INTs, we examined the relationship between INTs and the aperiodic (1/f) components of the EEG power spectrum. The aperiodic exponent is often interpreted as an index of E/I balance, with changes in slope reflecting the interaction between excitatory and inhibitory activity^16,54,55^. Meanwhile, the offset, is thought to reflect the level of aggregate neural spiking^16,54,55^. Here, we found that both a lower aperiodic exponent (increased E/I balance) and a lower offset (reduced neural spiking) were significantly associated with shorter INTs, suggesting that cortical networks with heightened E/I balance and reduced spontaneous firing exhibit faster temporal dynamics. These results provide converging evidence that maturation of local circuitry, particularly increased inhibitory control leading to the reduction of spontaneous activity, contributes to the developmental tuning of temporal integration. This implies that during adolescent maturation, cortical networks become more selective and efficient, reducing the need for prolonged temporal integration while allotting larger windows for cognition and favoring more brief, transient responses, consistent with previous literature on transient events during task states in humans and monkeys^12,82^. Importantly, these results are consistent with known physiological changes in adolescence, such as the maturation of GABAergic interneurons, particularly PV+ fast spiking cells^36,77–81^, and the refinement of local synaptic architecture, such as synaptic pruning^23,43^. These interneurons are critical for producing synchronized neural activity and local gain control, which may lead to the necessary temporal precision for shorter integration windows. These findings provide evidence for circuit level mechanisms impacting macroscale temporal dynamics, allowing us to link how developmental shifts in local cortical physiology drive the refinement of neuronal processing.

In addition to physiological mechanisms, we investigated structural contributions to INT development by examining the relationship between INTs and intracortical myelin, as measured from 7T quantitative MRI. Myelination through adolescence^9,10^ increases the conduction velocity of action potentials and signal fidelity, enabling more synchronized and temporally precise integration across networks, necessitating shorter INTs^83–85^. Taken together, structural changes in pruning and myelination, along with the maturation of inhibitory circuitry and improved cortical SNR, contribute to a more balanced E/I circuitry. As the E/I circuitry becomes more balanced, the maturing inhibitory network, specifically from PV+ interneurons, improves the temporal precision of cortical firing by suppressing excessive or asynchronous excitation^36^. This shift promotes shorter INTs, particularly in association regions like the prefrontal cortex, which would support developmental improvements in complex information processes underlying the maturation of executive functions. Deep layer myelination in frontal and parietal regions did not show a significant main effect on INTs. However, a significant age-by-layer interaction indicates that developmental changes in deep layer myelination are linked to shorter INTs over adolescence. Importantly, this association was specific to INTs measured during resting state, and was not observed during task state, suggesting that myelination contributes to the baseline organization of INTs across development, rather than influencing their dynamic modulation during cognitive demands. In contrast, intracortical myelination in superficial layers, which provides intracortical connectivity and has been found to show persistent increases through adulthood^10^, was not associated with age related changes in INT. The association with deep layers, as opposed to superficial, may suggest that deep layer circuitry, known to support long-range connectivity to the subcortex and feedback processing^86^ and shows stabilization in adulthood^10^, may play a greater role in shaping spontaneous resting state INTs into adulthood, as opposed to task evoked dynamics. Furthermore, rapid dynamic changes in neural firing may be more relevant to deep layers that produce temporally precise outputs^87,88^, compared to superficial layers that leverage more persistent and recurrent activity^87,89^. Taken together, these findings suggest structural refinement may constrain the temporal window of local populations, promoting a shift toward faster temporal integration across adolescence.

Given that timescales exhibit a hierarchical organization corresponding to behaviorally relevant demands^17,27^, and that they are modulated by cognitive state^27^, we sought to investigate how INTs measured during rest or task would be associated with performance in a working memory task. Overall, we show that shorter INTs were associated with improved working memory accuracy as well as less response latency variability. Importantly, this relationship was particularly prominent in adolescence, where adolescents with shorter INTs in frontal and parietal regions demonstrated relatively more accurate memory guided saccades, indicating a functional advantage of reduced temporal integration windows on behavior. Notably, this advantage was not observed in adults, suggesting that INT reductions across adolescence may be sufficient to support task performance in adulthood. In contrast, adolescents may require more dynamic upregulation, or task-related increases, in INTs to achieve comparable performance, indicating that INT modulation may serve a compensatory role during this developmental window. Previous work has linked INTs to various aspects of cognition, including working memory^32^, but to our knowledge, this is the first study to demonstrate that INTs predict cognitive performance across adolescence.

Together, our findings position INTs as a as a key signature of adolescent temporal reorganization that supports increasingly flexible and efficient information processing. By leveraging a multimodal, cross species dataset, including scalp EEG, intracranial recordings from humans and nonhuman primates, structural MRI, and behavior, we demonstrate that INTs shorten across adolescence in a manner shaped by the underlying circuit physiology and structural maturation. Additionally, these temporal refinements coincide with improvements in working memory, suggesting that adolescence is not only a period of significant structural and functional maturation, but also one of temporal reorganization in which windows of integration are dynamically modulated to meet the demands of external inputs. While this study offers novel insights into the developmental refinement of INTs, several limitations should be noted. First, EEG-derived INTs reflect large scale, surface level dynamics that are influenced by volume conduction. Although convergence with sEEG and macaque LFPs seeks to address this gap, the indirect nature of EEG are indirect. Second, our sEEG dataset is a small sample and was only collected during resting state, limiting our ability to bridge surface level task dynamics to intracortical dynamics within the same species. We also acknowledge that the ACF and PSD are mathematically related. However, our goal here was not to examine this redundancy, but rather to test whether INTs, derived from the temporal domain, show associations with specific aspects of the aperiodic component (i.e. the exponent and offset). Altogether, our findings demonstrate that INTs capture developmental shifts in neural signal dynamics across species and modalities, providing a mechanistic framework for understanding how cortical maturation supports increasingly specialized information processing in adolescence.

## Methods

### Developmental dataset

#### Participants

This study analyzes data from a healthy, neurotypical, accelerated longitudinal cohort, recruited from the greater Pittsburgh area. Data was collected on 164 participants (87 assigned female at birth), between 10-32 years of age. Participants were recruited as part of an accelerated cohort design with up to 3 visits at approximately 18mo intervals, for a total of 347 visits. Each time point consisted of three sessions: a behavioral (in-lab) session, a 7T MRI scan, and an EEG session, typically occurring on different days within 1-2 weeks. Participants were excluded if they had a history of loss of consciousness due to a head injury, non-correctable vision problems, learning disabilities, a history of substance abuse, or a history of major psychiatric or neurologic conditions in themselves or a first-degree relative. Patients were also excluded if any MRI contradictions were reported, including but not limited to, non-removable metal in their body. Following data quality control and exclusion criteria, the final EEG dataset included data from 161 individuals and 290 longitudinal sessions (70 individuals with 1 session, 53 individuals with 3 sessions, and 38 individuals with 3 sessions). Participants or the parents of minors gave informed consent with those under 18 years of age providing assent. Participants received payment for their participation. All experimental procedures were approved by the University of Pittsburgh Institutional Review Board and complied with the Code of Ethics of the World Medical Association (Declaration of Helsinki, 1964).

### Memory guided saccade task (MGS)

During the EEG session (see acquisition details below), participants completed a memory-guided saccade (MGS) task designed to assess working memory^12^. Each trial began with a 1-second fixation on a central blue cross, followed by the presentation of a peripheral cue embedded within a scene image (for a separate study), appearing at an unknown horizontal location (either 12.5° or 22.2° to the left or right of center). Participants executed a visually guided saccade (VGS) to the cue and maintained fixation for 1 sec. After the cue disappeared, they returned gaze to central fixation and maintained it through a variable delay period (6–10 seconds), during which they retained the cue location in working memory. When the central fixation cross was extinguished, participants performed a memory-guided saccade to the remembered target location without any visual cues. Trials concluded with a white fixation cross marking the inter-trial interval (1.5–15 seconds). Participants completed three runs of 20 trials each. A visual representation of the task can be seen in Supplemental figure 1.

Task performance was measured using horizontal electrooculogram (hEOG) signals recorded from facial electrodes. Prior to the task, participants underwent a calibration procedure involving sequential fixation on 20 dots spanning the horizontal midline. This calibration was used to generate a mapping between hEOG voltage and screen position. During the task, hEOG signals were aligned to stimulus triggers to calculate VGS and MGS latencies, defined as the time from recall onset to eye movement initiation, and saccadic accuracy, defined as the deviation between the MGS fixation point and the original VGS target location. The primary behavioral outputs from the MGS task were the average accuracy (measured as the degrees away from the intended target, averaged across trials), accuracy variability (the standard deviation of the accuracy measures across trials), latency (measured as the time difference between the beginning of the recall period and the initiation of the eye movement, averaged across trials), and the latency variability (the standard deviation of the latency measures across trials).

### Electrophysiological (EEG) acquisition and preprocessing

Concurrent electrooculogram (EOG) and high-impedance EEG were recorded using a 64-channel Biosemi ActiveTwo system in the PWPIC Psychophysiology Laboratory. Sessions took place in an electromagnetically shielded room, with stimuli presented on a computer approximately 80 cm from participants. Data was collected during resting state, across four alternating one-minute eyes-open and eyes-closed conditions, and the MGS task (described above). EEG data were initially sampled at 1024 Hz and later resampled to 150 Hz during preprocessing. Scalp electrodes were referenced to mastoid electrodes, chosen for their proximity to the scalp and low noise levels. A bandpass filter of 0.5–90 Hz was applied to control for low frequency drifts and muscle movement.

Preprocessing followed a modified EEGLAB^90^-compatible pipeline, which included the removal of flatline channels (≥8 seconds), low-frequency drifts, noisy channels (>5 SD from the mean), brief spontaneous bursts, and incomplete data segments. Removed channels were interpolated using neighboring electrodes, and the data were re-referenced to the average. Independent component analysis (ICA) was then used to identify and remove eye-blink artifacts. To account for line noise, we filter for 60hz. This preprocessing pipeline has been previously described in McKeon 2023, McKeon, 2024a, and McKeon, 2024b^12,13,16^.

### Intrinsic Neural Timescale Estimation via the autocorrelation window

#### Autocorrelation Window

The current study employed ACW-50 to quantity INTs on all 64 electrodes for all participants. Previous work has shown shorter INTs (as measured by the ACW) in unimodal sensory regions compared to higher order transmodal regions^17,22^. Thus, to assess differences in the ACW between regions electrodes were grouped into three broad areas: frontal (electrodes Fp1, AF1, AF5, F7, F5, F3, F1, FC1, FC3, FC5, FT9, C1, Fp2, AFz, AF2, AF6, F8, F6, F4, F2, Fz, FCz, FC2, FC4, FC6, FT10, C2, Cz), parietal (electrodes P9, P7, P5, P3, P1, PO3, PO7, PO9, POz, Pz, P10, P8, P6, P4, P2, PO4, PO8, PO10), and occipital (electrodes O1, I1, Oz, O2, I2). ACW was calculated on every trial from the MGS task in 2 second chunks of data across the variable delay period lengths (6, 8, or 10 seconds) and then averaged across trials. We then compared the ACW derived from the 2-second windows across the variable delay periods and found no significant differences (Supplement Figure 2). Thus, the ACW values from the 2-second windows were averaged together to create one measure per trial. Trial level values were then averaged to create a per subject measure of ACW. For consistency, the resting state data was also broken up into 2-second data chunks. The ACW from these 2 seconds windows of data were averaged together to get one measure. The autocorrelation window (ACW) was derived from the autocorrelation function (ACF), which quantifies the similarity of a signal with itself at different time lags. The time lag when the ACF delays to 50% of its maximum was calculated and deemed ACW-50 (as this is the only ACW measure we report in this study; we will further refer to it as simply the ACW). This represents the temporal “half-life” of the signal’s self-similarity and has been previous reported in the literature^21,24^. A threshold of 2 standard deviations from the mean was used to exclude statistical outliers.

#### Associations between working memory vs resting state

Previous work has shown that INTs are modulated by task state^27^, thus we sought to assess whether the ACW in frontal, parietal, and/or occipital regions would vary between resting state and various working memory states (i.e. delay vs fixation periods). To do so, we implemented two-tailed paired sample t-tests on the ACW from the delay period of our MGS task vs resting state. We repeat this assessment between MGS delay and fixation to interrogate whether different aspects of the same task elicit different responses.

### Developmental analyses

To assess developmental changes in the ACW across adolescence, we employed semiparametric GAMMs (generalized additive mixed models) with penalized splines using the mgcv package in R^91^. Independent GAMMs were run for each regional analysis (frontal, parietal, and occipital). The developmental models were fit with ACW as the dependent variable modelling age as a smooth term. Fixed effects for sex were also included in the model, as well as a random intercept per participant. To prevent overfitting, a maximum basis complexity (k) of 3 was used. Significance values were Bonferroni corrected (n=3). In addition to the main development models, independent GAMMs per region with an age-by-task interaction term were fit to assess whether age-related patterns differed across task states. These GAMMs were fit with ACW as the dependent variable, smooth terms for age, a factor-smooth interaction term between age and task, as well as a random intercept per participant. Fixed effects of sex were also included in the model.

We then sought to assess the rate of ACW change with age in each of region. A GAMM (generalized additive mixed model) model was fit with the ACW as the dependent variable, age as a smooth term, controlling for sex, and with a random intercept per participant. The first derivative of the age spline was computed to estimate the rate of developmental change using the gratia package in R^92^, using 200 equally spaced age intervals. To index the average rate of ACW change across the age spline, the average first derivative was computed, where a more negative average derivative is indicative of greater maturational decline.

In addition to assessing ACW in task vs resting state, we sought to characterize whether ACW effects were driven by the adults or adolescents within our sample. To do so, we implemented two-sided paired sample t-tests on the ACW from the delay period of our MGS task vs resting state for both adults (>= 18) and adolescence (< 18).

### Associations with working memory performance

During EEG data collection, participants completed a memory guided saccade task (described above). The primary behavioral outputs from the MGS task were the average accuracy, accuracy variability, latency, and latency variability. To assess the effects of the ACW within each region on working memory performance, we used GAM models, independently run for each performance metric. We first tested for significant main effects, with the behavioral measure as the dependent variable, age as a smooth term, fixed effects of ACW, task state (resting or task), and sex. Participant specific random intercepts were modeled using a random effect term to account for repeated measures. In addition to the main effect models, independent GAMs per region with an ACW-by-task interaction term were fit to assess whether ACW-related patterns on performance differed across task states. These GAMs were fit independently with each behavioral measure as the dependent variable, smooth terms for age, an interaction smooth term between ACW and task, as well as a random intercept per participant. Fixed effects of sex and task state were also included in the model. For each behavioral measure, p-values were Bonferroni corrected based on the 3 cortical regions of interest.

Additional analyses were conducted on the associations that yielded significant results to further interrogate how the relationship between ACW and behavior changes across development. To do so we modeled the age-varying effect of ACW on each behavioral measure using a semiparametric GAM framework. This is conceptually equivalent to a time-varying effect model (TVEM), allowing the strength of the ACW-behavior relationship to change across age. Specifically, we modeled behavior as a function of smooth age terms, with an interaction smooth term between age and ACW. The model also included fixed effects for task state and sex, as well as a participant-specific random intercept. To characterize the age-varying effect of the ACW-behavior association, predicted values were visualized using ggpredict in R. The first derivative of the age smooth was computed to find the significant points of developmental change, identified as the age ranges where the derivative confidence interval did not cross zero.

### Associations with intracortical myelin

#### MP2RAGE acquisition

MRI data acquisition and preprocessing have been previously described in Sydnor et al., 2025^10^. Briefly, MRI data were collected on a 7T Siemens Magnetom (MR B17) using parallel transmission (pTx) to enhance B1+ homogeneity^93^. A magnetization prepared 2 rapid acquisition gradient echoes (MP2RAGE) sequence was used to generate T1-weighted (UNI) images and quantitative T1 maps. Two GRE images, acquired at different inversion times to reduce B1-, proton density, and T2* effects, were combined to generate the MP2RAGE. Data were acquired with a GRAPPA acceleration factor of 4, inversion times of 800 ms (INV1) and 2700 ms (INV2), RT of 6000ms, echo time of 2.87 ms, 4 (INV1) and 4 (INV2) degree flip angles, and 1 mm isotropic voxel resolution. UNI images were denoised using a regularized combination of the inversion images^94^.

#### MP2RAGE-based cortical myelin imaging

Quantitative T1 maps, derived from the MP2RAGE sequence, were used to calculate the longitudinal relaxation rates (R1=1/T1), a validated proxy for cortical myelin content, as detailed in Sydnor et al., 2025^10^. Briefly, T1 maps were corrected for B1+ inhomogeneity using the UNICORT (unified segmentation based correction of R1 brain maps for transmit field inhomogeneities) algorithm. Person-specific cortical surfaces were then generated from UNI images using FreeSurfer’s longitudinal processing stream (version 7.4.1). Volumetric R1 maps were calculated from quantitative T1 images using 3dcalc from AFNI (R1 = 1/T1 in sec^-1^) and were resampled onto each participant’s cortical surface using FreeSurfer volume-to-surface projection. R1 was sampled at 11 cortical depths between the pial (0% cortical thickness) and gray-white (100% cortical thickness) boundary, in 10% increments. Using the cortex volume fraction, which quantifies the fraction of the signal that comes from gray matter as opposed to white matter, we included 7 cortical depths where the average cortex volume was >90%. We then divided these cortical depths into superficial depths (20-40% of cortical thickness) and deep depths (50-80% of cortical thickness). All R1 data used in the present study were obtained directly from this prior work and were not reprocessed.

#### Atlas of EEG electrode positions

In order to relate cortical R1 to EEG ACW, R1 values were quantified in cortical locations proximal to EEG electrodes. This required the creation of a surface-based atlas of EEG electrode positions by projecting the electrode positions from the scalp to the brain’s surface using Brainstorm version 03^95^. Details about the creation of the EEG surface atlas can be found in Sydnor et al. 2025^10^. Briefly, cartesian coordinates of all 64 electrodes were uploaded in Brainstorm, placed on the scalp of a generic head model, and projected onto the cortical surface of an MNI152 template. Labels were then projected to fsaverage surface using neuromaps and then transformed from the fsaverage template to participant-specific cortical surfaces from FreeSurfer. R1 was then averaged into superficial depths and deep depths in regions corresponding to each electrode label. Labels corresponding to frontal, parietal, and occipital regions were averaged together in order to relate R1 measures in those regions to EEG ACW measures.

#### Modeling R1 associations with ACW

To study the associations between cortical R1 and EEG-derived ACW we leveraged developmental GAMs with R1 (superficial or deep) as the dependent variable, age as a smooth term, ACW as a smooth term, and a random intercept per participant. GAMMs were fit separately for the 3 cortical regions (frontal, parietal, and occipital) for task and resting state ACW values. To assess the strength and significance of the associations between R1 and the ACW in superficial and deep layers, extracted the F-statistic and corresponding p-value for the ACW smooth term from each model. For task and resting state conditions, p-values were Bonferroni corrected based on the 3 cortical regions of interest.

We then sought to interrogate for differences in the strength of the R1-ACW relationship by depth. Thus, we fit GAMMs with a depth interaction term, separately for the task and resting state conditions, as well as, for frontal, parietal, and occipital regions. The models included R1 (superficial or deep) as the dependent variable, age as a smooth term, and an ordered factor smooth interaction between depth and ACW. Depth was also included as a linear covariate along with a random intercept per participant.

### sEEG Dataset

#### Participants

sEEG recordings were performed in 14 neurosurgical patients with drug-resistant epilepsy as part of clinical evaluation for epilepsy surgery. See Supplement Table 1 for patient-participant characteristics. Written consent and assent were obtained from all participants. The research protocol was approved by the University of Pittsburgh Institutional Review Board (STUDY19010005).

#### Acquisition and preprocessing

Each patient underwent implantation of depth electrodes as previously described Abel et. al 2018^96^. Recording sites were selected based on clinical necessity for seizure localization. Briefly, Dixi Medical Microdeep electrodes were employed, featuring a diameter of 0.8 mm, a contact length of 2 mm, and a center-to-center spacing of 3.5 mm. Each electrode comprised between 8 and 18 contacts, which are referred to as channels. Neural recordings were collected during 5-minute eyes open resting state, recorded at 1kHz using the Ripple Grapevine Nomad system (model R02000) with real-time notch filtering applied at 60, 120, and 180 Hz. Data was resampled to 512 Hz and a common average reference (CAR) filter was used to remove noise shared across channels.

#### Autocorrelation Window Calculation

The autocorrelation window (ACW) was calculated using the same approach described previously. In brief, resting-state recordings from each channel were segmented into 2-second epochs. For each epoch, the autocorrelation function (ACF)—which quantifies the similarity of a signal with itself across varying time lags—was computed. The ACW was defined as the time lag at which the ACF decayed to 50% of its maximum value, a measure referred to as ACW-50. As this is the only ACW metric reported in the current study, it will be referred to hereafter simply as the ACW. This metric represents the temporal “half-life” of the signal’s self-similarity and has been previously described in the literature^21,24^. Only channels in non-seizure sites, determined by x, were used in the main analysis. A threshold of 2 standard deviations from the mean was used to exclude statistical outliers.

#### Developmental analyses

As in our longitudinal developmental cohort, we assessed age-related changes in ACW across adolescence using semiparametric GAMMs (generalized additive mixed models) with penalized splines using the mgcv package in R^91^. Independent GAMMs were run for each regional analysis (frontal, parietal, and occipital). The developmental models were fit with ACW as the dependent variable, age as a smooth term. Fixed effects for a random intercept per participant. To prevent overfitting, a maximum basis complexity (k) of 3 was used. Significance values were Bonferroni corrected (n=3). In addition to the main development models, independent GAMMs per region with an age-by-task interaction term were fit to assess whether age-related patterns differed across task states. These GAMMs were fit with ACW as the dependent variable, smooth terms for age, an interaction smooth term between age and task, as well as a random intercept per participant.

### Non-human primate dataset

#### Subjects

Neurophysiological data were collected from 8 rhesus monkeys (Macaca mulatta), comprised of 6 males and 2 females. All surgical procedures and animal use protocols were reviewed and approved by the Institutional Animal Care and Use Committees at Wake Forest University and Vanderbilt University, in compliance with the U.S. Public Health Service Policy on Humane Care and Use of Laboratory Animals and the National Research Council’s Guide for the Care and Use of Laboratory Animals.

#### Developmental Stage

Developmental measures of the macaques were tracked on a quarterly basis before, during, and after neurophysiological recordings. Bone maturation via X-rays of upper and lower extremities were used to determine each monkey’s developmental process. Skeletal assessment was used to best capture physical development^97–99^ and individually aligned to growth trajectories for each macaque. The time of each monkey’s distal tibial epiphyseal closure was used to define a mid-adolescence age for each monkey, as observed by veterinary professionals evaluating the X-rays, blind to findings of other aspects of the study.

#### Oculomotor delayed response (ODR) Task

Monkeys were trained to perform an oculomotor delayed response (ODR) task similar to the MGS task used in our EEG study^100^ (Figure 5D), a spatial working memory paradigm in which subjects were required to remember the location of a briefly presented cue. The cue—a 1° white square—was displayed for 0.5 seconds at one of eight possible locations arranged equidistantly (45° apart) on a 10° eccentric circle. Following a 1.5 or 3-second delay period, the fixation point was extinguished, signaling the monkey to make a saccade to the remembered cue location within 0.6 seconds. A correct response required the saccade to land within a 6° radius window centered on the cue location and to be maintained for at least 0.1 seconds. Correct trials were reinforced with liquid rewards, typically fruit juice. Eye position was monitored using an infrared eye-tracking system (ISCAN RK-716; ISCAN, Burlington, MA) with a sampling rate of 240 Hz.

#### Surgery and Neurophysiology

Following initial attainment of asymptotic performance on the behavioral tasks, monkeys underwent surgical implantation of a 20-mm diameter recording cylinder over the prefrontal cortex. The placement of the recording chamber and electrode penetrations was guided by structural MRI and co-registered using the BrainSight system (Rogue Research, Montreal, Canada).

Neurophysiological recordings in areas 8a and 46 of the dorsolateral prefrontal cortex were performed at intervals of approximately three months, spanning ages 3.4 to 6.2 years, using either glass- or epoxylite-coated tungsten microelectrodes (FHC, Bowdoin, ME), with a shaft diameter of 250 μm and impedance of 4 MΩ at 1 kHz. Neural signals were amplified, band-pass filtered between 500 Hz and 8 kHz, and digitized at a temporal resolution of 25 μs using a modular acquisition system (APM system, FHC, Bowdoin, ME). On average, each time point for each animal included 19 behavioral sessions. Between time points, animals were returned to their home colony and did not receive any further training or task exposure until the next scheduled session.

#### Autocorrelation Window Calculation

The autocorrelation window (ACW) was calculated using the same approach described previously. In brief, recordings from each channel, for each behavioral session, for every trial, for each monkey, were separated into the delay and fixation epochs of the ODR task. For each epoch, the autocorrelation function (ACF)—which quantifies the similarity of a signal with itself across varying time lags—was computed. The ACW was defined as above, as the time lag at which the ACF decayed to 50% of its maximum value, a measure referred to as ACW-50. As this is the only ACW metric reported in the current study, it will be referred to hereafter simply as the ACW. This metric represents the temporal “half-life” of the signal’s self-similarity and has been previously described in the literature^21,24^. A threshold of 2 standard deviations from the mean was used to exclude statistical outliers.

#### Developmental modeling

To examine developmental changes in the autocorrelation window during task performance, we fit a generalized additive mixed model (GAMM) using the bam() function (gam for large datasets) from the mgcv package in R^101^. The model included smooth terms for age nested within individual monkeys to capture subject-specific developmental trajectories, as well as separate smooths for behavioral session, trial number, and channel-by-monkey interactions to account for within-session and within-subject variability. Additional random effects were included for neuron recording area, monkey identity, and recording epoch (delay or fix). A smooth interaction between age and epoch was also included to model age-related effects that varied across experimental epochs.

To visualize developmental trajectories of ACW across adolescence, we generated smooth fits using generalized additive mixed models (GAMMs) with ggplot2. Individual trajectories were plotted for each monkey using subject-specific smooths with fixed basis complexity (k = 3), providing within-subject developmental trends. A group-level trend line was overlaid using a GAM with an unfixed smoothing parameter, capturing the overall developmental pattern across the cohort. Line color indicated monkey identity, while line type distinguished between experimental epochs. Shaded confidence intervals were included for both individual and group fits.

## Data availability

Data will be made available upon request

## Code availability

All original code generated for this study is available on github at https://github.com/LabNeuroCogDevel/SignalComplexityAcrossAdolescence. In addition, a detailed guide to code implementation describing all analytic steps and statistical analyses is provided along with the github repository at https://labneurocogdevel.github.io/SignalComplexityAcrossAdolescence/.

## Acknowledgements

We thank the University of Pittsburgh Clinical and Translational Science Institute (CTSI) for support in recruiting participants, as well as their support by the National Institutes of Health through grant number <u>UL1TR001857</u>. We thank Matthew Missar, Laurie Thompson, and Vivian Lallo for their work involving our data collection.

