## Supplement for "Cross-species evidence for the refinement of intrinsic neural timescales supporting executive system maturation through adolescence"

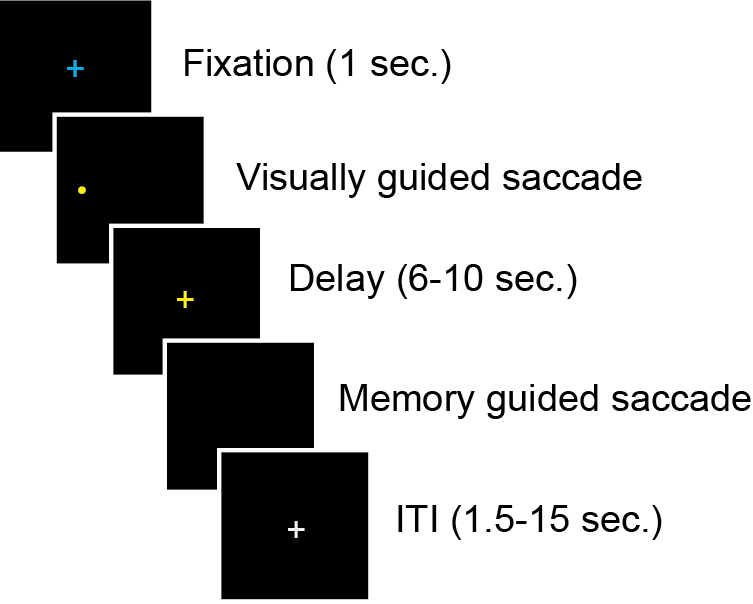

Supplement Figure 1. Visual representation of the MGS task.

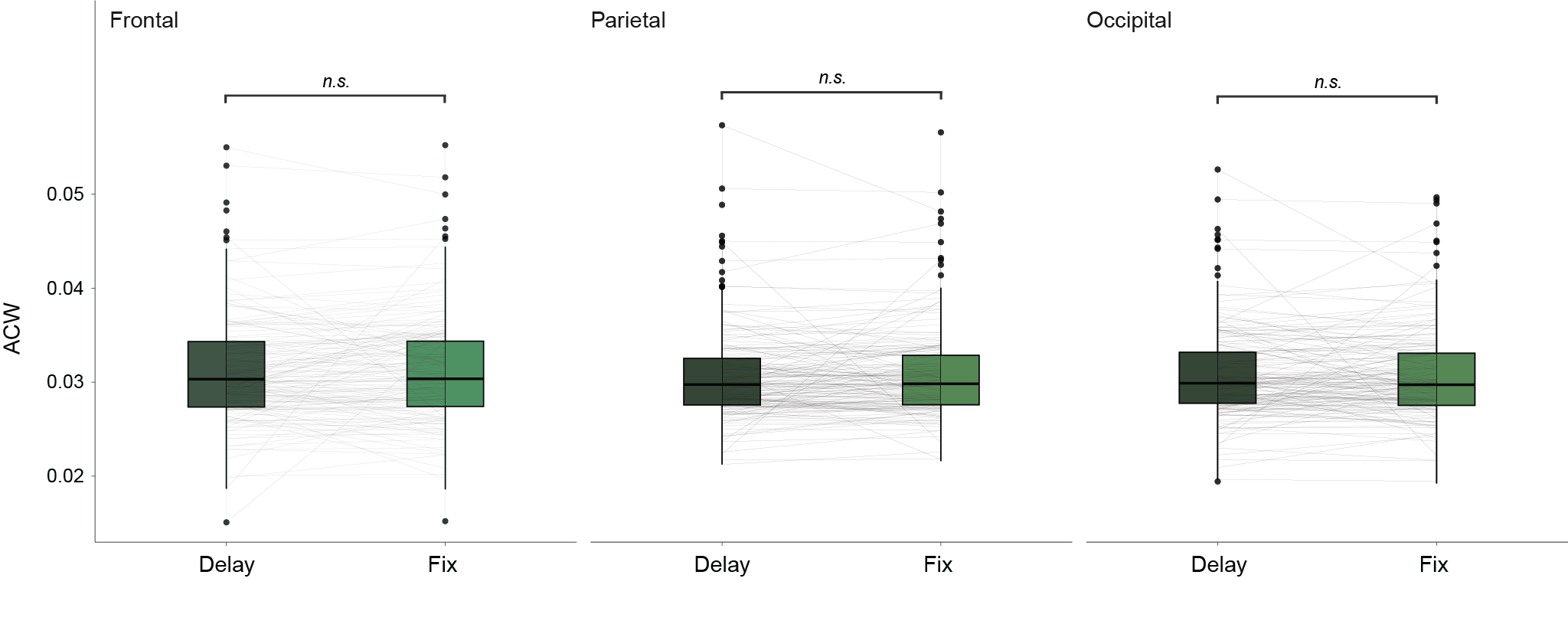

Supplement Figure 2. INTs reflect a cognitively engaged state. Boxplots compare the autocorrelation window between MGS delay period and fixation period across all participants in the frontal, parietal, and occipital lobes. No significant differences were found between the two epochs, allowing us to average the ACW value of the two epoch together into one ‘task state’.

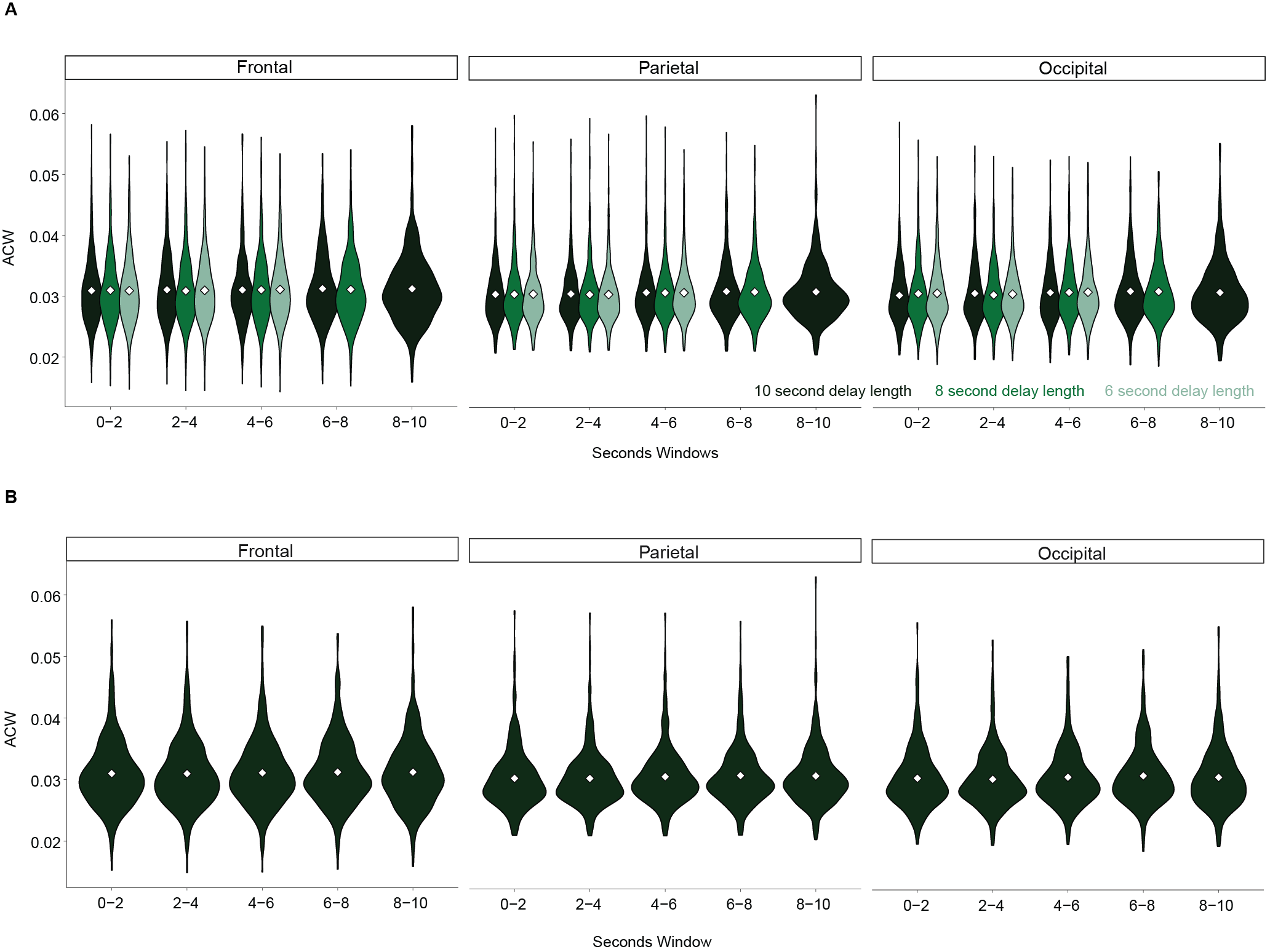

Supplement Figure 3. Average ACW for the three delay period lengths (10, 8 or 6 seconds) broken up into 2 second windows. Violin plots show the distribution of the data. White diamonds represent the mean. (A) Analysis of the ACW_50 across different epochs and time windows in the frontal, parietal, and occipital regions. A generalized additive model (GAM) was used on each region to assess the influence of epoch (Delay6, Delay8, Delay10), and time window (seconds window) on the ACW while controlling for age and sex. No significant differences were found in any of the three regions between the epochs within any specific time window, as all epoch × seconds window interactions were non-significant (all p > 0.05). These findings suggest that ACW_50 values are largely stable across epochs, with minimal influence from the specific time window in the frontal, parietal nor occipital regions. (B) Based on the previous analysis, we averaged the ACW from the three different delay period lengths for each two second window. A generalized additive model (GAM) was used on each region to assess the influence of time window (seconds window) on the ACW while controlling for age and sex. No significant differences were found in any of the three regions across the specific time windows (all p > 0.05). These findings suggest that ACW values are largely stable across the delay period in the frontal, parietal nor occipital regions. This led us to average the ACW from each two second window resulting in an average ACW value for each subject’s delay period.

Supplemental Table 1. sEEG subject demographics

|  | Age (years) | Sex | WASI-II Score | Number of non-epileptic channels/ all |
| --- | --- | --- | --- | --- |
| Summary | 15.4 (3.5) | 71% Male | 92 (18) | 57/150 |
| P1 | 18 | Male | 95 | 32/127 |
| P2 | 15 | Male | 81 | 71/117 |
| P3 | 9 | Male | 113 | 69/116 |
| P4 | 15 | Female | 102 | 61/104 |
| P5 | 18 | Male | 94 | 18/101 |
| P6 | 12 | Female | 85 | 62/128 |
| P7 | 9 | Male | 67 | 25/120 |
| P8 | 21 | Female | 98 | 58/199 |
| P9 | 17 | Male | 81 | 92/164 |
| P10 | 18 | Male | 60 | 81/205 |
| P11 | 16 | Male | 82 | 62/256 |
| P12 | 19 | Male | 117 | 38/162 |
| P13 | 13 | Female | 125 | 71/172 |
| P14 | 16 | Male | 88 | 67/130 |
